# Light environment drives evolution of color vision genes in butterflies and moths

**DOI:** 10.1101/2020.02.29.965335

**Authors:** Yash Sondhi, Emily A. Ellis, Jamie C. Theobald, Akito Y. Kawahara

## Abstract

Opsins are the primary light-sensing molecules in animals. Opsins have peak sensitivities to specific wavelengths which allows for color discrimination. The opsin protein family has undergone duplications and losses, dynamically expanding and contracting the number of opsins, throughout invertebrate evolution, but it is unclear what drives this diversity. Light availability, however, appears to play a significant role. Dim environments are associated with low opsin diversity in deep-sea fishes and cave-dwelling animals. Correlations between high opsin diversity and bright environments, however, are tenuous. Insects are a good system to test whether opsin expansion is associated with greater light availability because they are enormously diverse and consequently display large variation in diel activity. To test this, we used 200 insect transcriptomes and examined the patterns of opsin diversity associated with diel-niche. We focused on the butterflies and moths (Lepidoptera) because this group has significant variation in diel-niche, substantial opsin recovery (n=100), and particularly well-curated transcriptomes. We identified opsin duplications using ancestral state reconstruction and examined rates of opsin evolution, and compared them across diel-niches. We find Lepidoptera species active in high light environments have independently expanded their opsins at least 10 times. Opsins from diurnal taxa also evolve faster; 13 amino acids were identified across different opsins that were under diversifying selection. Structural models reveal that four of these amino acids overlap with opsin color-tuning regions. By parsing nocturnal and diurnal switches, we show that light environment can influence gene diversity, selection, and protein structure of opsins in Lepidoptera.

## Introduction

Opsins are the primary light sensing molecule in animals. Lepidoptera has over 150,000 species with large variation in the activity time and light environment (diel-niche). Opsin diversity has yet to be studied comprehensively across the entire order and multiple diel-niches. Here we provide a novel link between visual genotype and light environment phenotype. We analyze a large dataset of 200 insect transcriptomes to study opsin diversity and its association with diel-niche. By comparing opsin diversity across 100 nocturnal and diurnal lepidopterans, we find that opsin diversity increases in bright light environments. Bright light environments are also associated with higher rates of opsin evolution, and some of these rapidly evolving sites are implicated in tuning vision.

Vision is a fast and reliable sensory modality, useful to detect shape, color and distance of signals. Eyes are diverse, convergently evolved structures that detect the direction and intensity of light in an animal’s environment^1^. Although eyes are often considered inseparable from vision, light sensitivity likely predates the origin of eyes^2^. Bacteria, fungal spores and dinoflagellates lack identifiable eyes, but can still detect and respond to light^3–5^.

The visual pigment rhodopsin is the primary light sensing molecule in all animals ranging from insects to primates. Rhodopsin, or opsin, transduces light into a biologically meaningful signal by activating a G-Protein and initiating the phototransduction signaling cascade. Large-scale phylogenomic analyses have found many duplications and losses — which we refer to as expansion and contraction — in the opsin protein family across invertebrates^6,7^. The water flea, *Daphnia pulex* possesses 46 opsins^8^, dragonflies have 12-15^9^ and mantis shrimp have 12-33^10^. However, any more than four spectral channels offer diminishing returns in extracting information from natural scenes^11,12^, so why do some animals have so many opsins?

Opsins may expand and contract for non-adaptive reasons. Non-adaptive forces usually cause random sequence evolution, and unless the duplicated opsins are co-opted for visual use, they usually become pseudogenes. Adaptive evolutionary forces are more likely to cause consistent, repeated, and persistent patterns of opsin diversification, such as with mate choice in guppies and butterflies^13,14^, flower foraging in bees and wasps^15^ and light intensity environment with many nocturnal animals^16,17^.

Sensory modalities such as smell, electromagnetic reception and touch are more reliable than vision in dim environments. Resource allocation trade-offs can cause loss of stabilizing selection on inefficient sensory systems and can cause a downregulation of opsins, leading to them becoming pseudogenes, a pattern seen in nocturnal mammals^18^, cave-dwelling crayfish^19^, and deep-sea organisms^20–22^. If diminished light availability causes reduced opsin expression and loss, abundant light, conversely, may cause higher opsin expression, duplications, and diversification. A comparison of visual genes between diurnal and nocturnal Lepidoptera revealed elevated opsin expression in the diurnal species^23^. Similarly, a study of opsin evolution across fireflies revealed higher amino acid transition rates in diurnal fireflies, across 4 independent diel switches^24^.

Lepidoptera opsin diversity has been studied in handful of model taxa — mostly diurnal butterflies^23,25,26^ — but opsin diversity has yet to be studied comprehensively across the entire order and multiple diel niches. Briscoe (2008) analysed the visual genes of 8 Lepidoptera species including two moths^25^, Xu et al. (2013) analysed 30 Lepidoptera including 12 moth species^25^, and Feuda et al. (2016) analysed 10 Lepidoptera species including four moths^28^. These studies annotate genomic opsins, and used gene trees instead of species trees for selection analyses, both of which can bias results. The few studies that examined opsin diversity and diel-niche association^27,28^ compare butterflies and moths, effectively using only a single diel switch. Lepidoptera have more than 100 recorded diel transitions^29^, and only by examining multiple independent diel-niche switches can we understand how light environment and diel-niche drive the evolution of their visual systems. To this end, we mine 176 Lepidoptera species transcriptomes for visual opsins and combine our annotations with publicly available data to present the most comprehensive survey of opsins across Lepidoptera. In parallel, we compiled diel-activity from the literature, natural history databases, and in consultation with experts.

## Results

### Opsin diversity patterns across insects

We examined patterns of opsin diversity across insects using assembled transcriptomes from InsectBase^30^ along with opsin annotations from Ensembl Metazoa ^31^ and previous studies^28^. Diel-niche assignment came from literature, and transcriptomes were annotated using a phylogenetically informed annotation approach (PIA)^32^. We recovered at least one opsin from more than half (28/50) of the insect transcriptomes, with a total of 79 opsin sequences. The final dataset included 45 insects (Table S2).

We reconfirmed the annotations using nucleotide and amino acid gene trees (Fig. S1, S2, supplementary information), and summarized opsin presence-absence and diel niche on a pruned insect tree (Fig. 1A). Our dataset of insect transcriptomes serves as a positive control for the annotation pipeline since the results agree with the broad patterns of opsin diversity published before^28^. However, the insect opsin dataset recovered opsins from too few nocturnal taxa to statistically compare opsin evolution between diel-niches. Therefore, we used similar methods on dataset of Lepidoptera with sufficient diel variation and better phylogenetic coverage to address this question.

**Figure 1:**
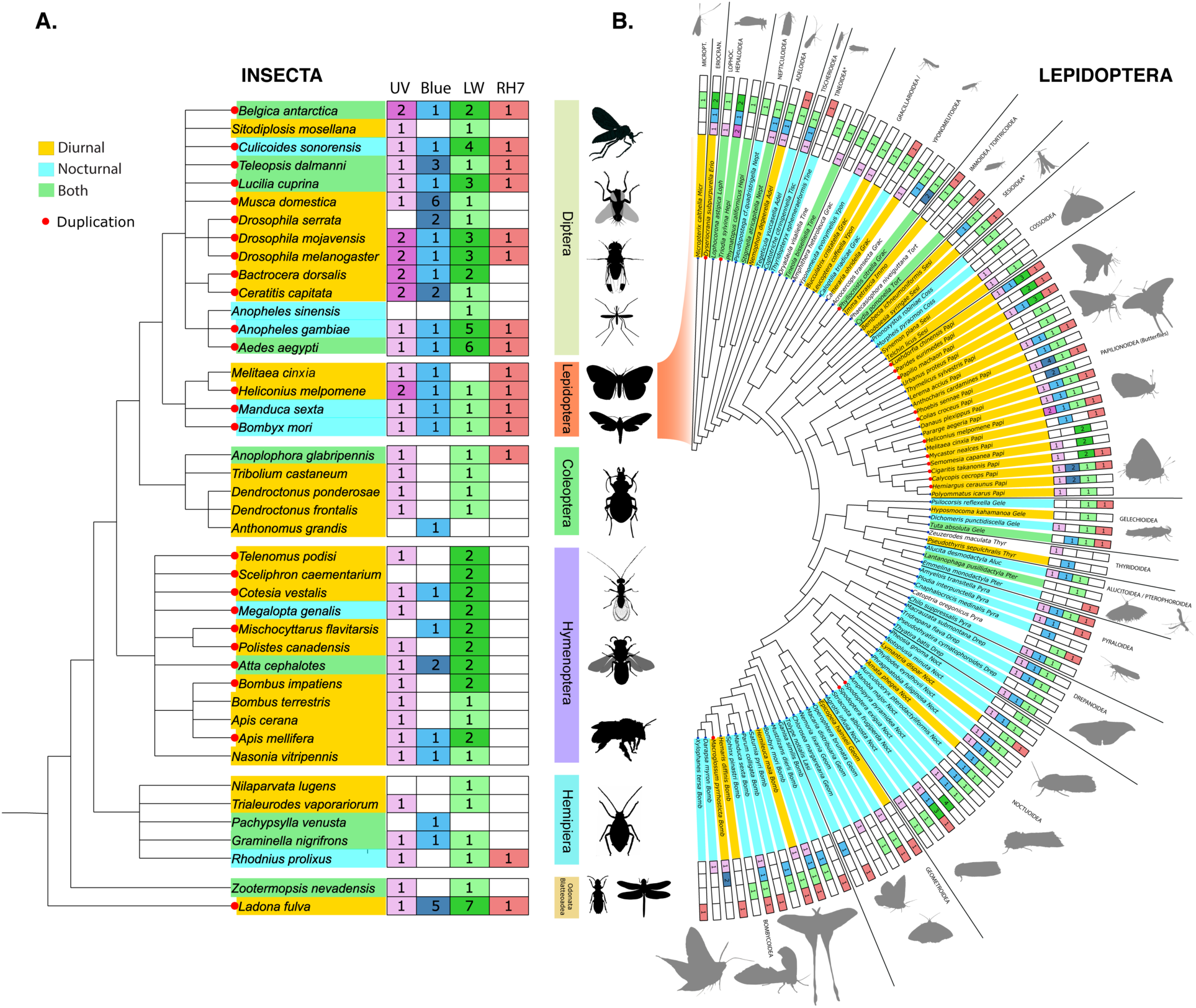
Opsin annotation and diel activity for Insecta and Lepidoptera. Taxa are color coded by diel-niche. Red dots at terminal nodes indicate duplications within a lineage for that species and darkened colors indicate a duplication in a particular opsin family. In the Lepidoptera tree, duplications (red dots) are more commonly associated with yellow and green colored taxa, which are at least partially active in bright light. In insects, data is sparse with too few nocturnal species (<25%) to establish such trends. A. Opsin annotation for 45 available insect transcriptomes and previously published literature. An NCBI taxonomy tree was grafted onto an accurate insect phylogeny obtained from Misof et. al 2014. Relationships at the level of order are based on the NCBI taxonomy tree and are only meant to be indicative of familial relationships and do not reflect current phylogenetic relationships between families. Insect silhouettes were taken from PhyloPic. B. Opsin annotation of 100 Lepidoptera species on a Lepidoptera phlyogeny. The tree is a pruned cladogram of tree of the most recent Lepidoptera phylogeny. (Kawahara et. al 2019).

### Lepidoptera opsin diversity and diel-niche association

We recovered at least one opsin from 114 of 175 Lepidoptera taxa, identified 265 opsin sequences (Table S1), and confirmed the opsin annotations by building gene trees (Fig. S1).

We mapped opsin diversity—specifically UV/RH4, Blue/RH5, LW/RH6 and RH7 opsin presence and absence—and diel-niche onto a well-resolved Lepidoptera phylogeny^33^ (Fig. 1B). We find that duplications occur more often in diurnal species than expected by chance (Table 1). Increased duplication in diurnal species was evident even after we excluded species with ambiguity in diel-niche assignment (Table 1).

**Table 1:** Chi-square values for nocturnal vs. diurnal Lepidoptera species, including and excluding crepuscular and “both” species. Duplications occur at a higher frequency in Diurnal species than expected by chance, even after accounting for various kinds of uncertainity in diel niche assignment. (All results are siginificant) D: diurnal, N: nocturnal #=(excluding uncertain diel states).

| Dataset | Chi-square coefficient | p-value | No. of samples<br>D:N | No. of duplications<br>D:N |
| --- | --- | --- | --- | --- |
| nocturnal vs. diurnal + crepuscular + both | 10.11 | 0.014 | 51:54 | 16:3 |
| nocturnal vs. diurnal + crepuscular + both<sup>#</sup> | 7.97 | 0.0047 | 51:46 | 16:3 |
| diurnal vs. nocturnal + crepuscular+ both | 9.32 | 0.0022 | 42:63 | 14:5 |
| diurnal vs. nocturnal + crepuscular + both<sup>#</sup> | 7.41 | 0.0064 | 42:55 | 14:5 |
| nocturnal vs. diurnal | 10.67 | 0.001 | 42:54 | 14:3 |
| nocturnal vs. diurnal* | 8.478 | 0.0035 | 42:46 | 14:3 |

We compared ancestral state reconstructions (ASR) of diel-niche and total number of opsins. We find that diurnal lineages and nodes have ancestrally higher opsin numbers (Fig. S3A). Opsin losses are notoriously difficult to confirm, since inferred losses could be due to poor opsin recovery or low-quality transcriptomes. To distinguish these from true losses, we performed an ASR for all four opsins (Fig. S3 A-D). If closely related lineages display a loss, ancestral nodes will show a high probability of loss (>50%); whereas, if losses are random, perhaps due to low quality data, then ancestral nodes will have 50% chance of loss.

#### Ultraviolet (UV)

50% (56/114) of the taxa recovered UV/RH4 opsins (Fig. 1, Fig. S3B). UV opsins had duplications in only 2 independent lineages; the diurnal *Heliconius melpomene*, in which UV opsin duplication has been recorded before^34^ and crepuscular *Triodia sylvina* (Hepialidae), an ancient lineage of ghost moths, known for swarming at dusk^35^. We did not find multiple lineages with a high probability of loss, but deeper nodes had a 50% chance of loss, likely indicating incomplete data, as opposed to true losses.

#### Blue

46.4% (53/114) of the taxa recovered Blue/RH5 opsins (Fig. 1, Fig. S1-3). Blue opsin duplications were present in 5 families and 7 genera, all diurnal species. The pierid *Phoebis sennae* and two lycaenid genera *Hemiargus* and *Polymmatus* also have blue opsin duplications. Behavioral and electrophysiological data have shown that these families, if not in these individual species, have functional duplications^25,36,37^. *Macrogolossum pyrrhosticta*, a diurnal sphingid hawkmoth, also had a blue duplication. Our ASR suggests true losses in four lineages in the Tortricidae, Crambidae, Noctuinae and Macroglossinae. For all other nodes, the chance of a loss was about 50%, likely indicating issues in opsin recovery and annotation.

#### Long Wavelength (LW)

83.3% (95/114) of taxa recovered LW/RH6 opsins, the highest of all three opsin families (Fig. 1, Fig. S3). LW opsins duplications were recorded in 8 families and 11 genera. Only two of these genera are nocturnal. We recovered previously reported LW duplications in one Lycaenidae, two Riodinidae, and three Papilionidae species^25^. Butterflies are among the few insects that can detect red, through LW duplication or filtering pigments^25,38^. Thus, any duplications might indicate true color expansion. Other diurnal or crepuscular species that had LW duplication were *Dyseriocrania subpurpurella* (Eriocraniidae) and *Triodia sylvina* (Hepialidae).

The tiger moth *Callimorpha dominula* and the castniid, *Paysandisia archon* – two diurnal moths not in this dataset – also have LW duplications, and *Paysandisia* at least can detect red pwavelengths ^28,36^. *Spodoptera—* of which some species are invasive pests—is nocturnal but has a red sensitive LW duplication. Many *Spodoptera* species are migratory, flying above the clouds in night skies may free them from low light constraints ^39,40^. *Tischeria quercitella*, a leaf mining moth, is the only other nocturnal species we found with a LW duplication, despite examining over 50 nocturnal species. Losses were identified in three distinct lineages, Geometrinae, Macroglossinae and, Smerinthinae; their presence was easier to confirm in LW than UV or Blue because of more complete LW recovery (Fig. S3D). Some Macroglossinae hawkmoths with a LW loss have a duplicated blue opsin, and electrophysiological investigation has shown that *Macroglossum stellatarum*, from the same clade, has three distinct sensitivity peaks^41^.

#### RH7

28% (33/114) of taxa recovered RH7 opsins with no duplications, consistent with other work (Feuda et al. 2016). Since RH7 is involved in circadian rhythm light sensing, it is possible that the time of sample collection affected the expression of opsins and might explain the lower recovery rate across our dataset.

### Opsin selection in Lepidoptera

We used PAML to estimate rates of selection (ω) and tested if these rates differed between nocturnal and diurnal Lepidoptera. For datasets that showed significant differences (ω), we used branch-site models in PAML to identify amino acids under selection. We tested if the analysis was sensitive to different sample sizes and starting trees (Fig. 2, Table 2, Table S4). To ensure that we compared opsin rates across the same group of species, we limited our analyses to species that recovered all three visual opsins. We were unable to use the entire dataset of recovered opsins for UV, Blue and Green opsins because the alignment included gaps, which could reduce the accuracy of the model. Since PAML requires stop codon free and unambiguous alignments, we removed sequences that created ambiguity in the alignment and trimmed the ends — not the middle, which confounds structural modelling — to ensure a clean alignment.

**Table 2:** ω (dN/dS) rates for null (M_0_) and branch models(M_2_) for Ultraviolet (UV), Blue, Long wavelength (LW) and RH7 opsins using species and gene trees and different number of taxa. Increasing the number of taxa shows significant differences between rates across nocturnal (ω_1_) and diurnal species (ω_2_) for color opsins but not for RH7 opsin, which served as a control. Using species tree shows differences for all three opsins. ω_0_: rates for null model, ω_1_: rates for diurnal species, ω_2_ for nocturnal species, n: number of taxa, LnL_0/2_ : log likelihood scores for M_0_/M_2_, df: degrees of freedom for M_2_,. 2LnL: twice the LnL_2_-LnL_0_, * highly significant p-value (<0.01)

| Opsin | $\omega_0$ | $\omega_1$ | $\omega_2$ | n | LnL<sub>0</sub> | LnL<sub>2</sub> | df | 2LnL | p-value |
| --- | --- | --- | --- | --- | --- | --- | --- | --- | --- |
| **M<sub>2</sub> vs. M<sub>0</sub> gene trees** |  |  |  |  |  |  |  |  |  |
| UV | 0.0328 | **0.0482** | **0.01643** | **24** | -13222.396 | -13196.06 | 48 | 52.67 | <0.001* |
| Blue | 0.03251 | 0.0403 | 0.02945 | 23 | -11310.806 | -11308.083 | 46 | 5.44 | 0.0196 |
| LW | 0.03264 | **0.0521** | **0.01173** | **24** | -11232.972 | -11174.84 | 48 | 116.27 | <0.001* |
| RH7 | 0.0653 | 0.0594 | 0.06719 | 33 | -22578.67 | -22577.68 | 66 | 1.978 | 0.15 |
| UV | 0.03058 | **0.0445** | **0.02324** | **14** | -6892.501 | -6887.289 | 28 | 9.6 | <0.001* |
| Blue | 0.0354 | 0.0343 | 0.03602 | 17 | -9822.971 | -9822.9126 | 34 | 0.115 | 0.735 |
| LW | 0.03893 | 0.0434 | 0.03445 | 17 | -6690.525 | -6689.4261 | 34 | 2.197 | 0.138 |
| **M<sub>2</sub> vs. M<sub>0</sub> species trees** |  |  |  |  |  |  |  |  |  |
| UV | 0.03206 | **0.0411** | **0.02609** | **24** | -13265.664 | -13260.007 | 49 | 11.314 | <0.001* |
| Blue | 0.03361 | **0.0465** | **0.02922** | **23** | -11333.887 | -11328.244 | 47 | 11.287 | <0.001* |
| LW | 0.03309 | **0.0495** | **0.02589** | **24** | -11247.362 | -11233.584 | 49 | 28.779 | <0.001* |

**Figure 2:**
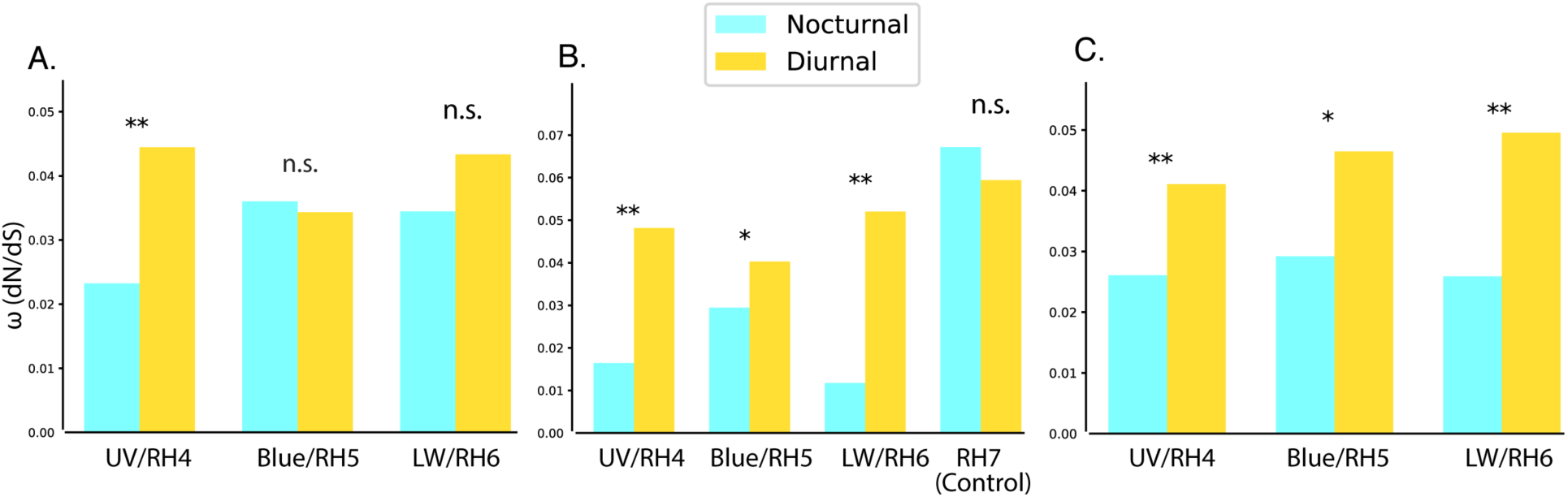
Opsin selection rates between nocturnal and diurnal Lepidoptera species for RH4/UV, RH5/Blue, RH6/LW and RH7 opsins. Diurnal and nocturnal species show differences in colour opsins for a few opsins, but on running the models with more sampling or species trees, these differences become more apparent, however RH7, a non-visual opsin, does not follow this trend. The test was two rates vs. 1 rate for the branch model vs. the null model with a Chi-square test. A. 14-17 taxon dataset run using opsin gene trees. B. 24 taxon dataset run using opsin gene trees. RH7 was analyzed using a larger dataset of 33 taxa. C. 24 taxon dataset run using species trees. * p-value <0.05, ** p-value <0.01

First, we ran PAML on a subset of the data (n=12–14)–of similar size as Feuda et al. (2016) and Xu et al. (2013)–to determine sample size effect, although taxon sampling across datasets was non-uniform. Of all three opsins, only UV opsin rates were significantly different between diel-niche for this dataset(Fig. 2A).

We next used a larger sample size and similar species across datasets (n=24–27). We found highly significant differences between diel-niche for UV and LW opsins (p-value <0.001) and moderately significant differences for Blue opsins (p-value <0.05). Each show relaxed selection in diurnal species. We used RH7 as a control because it is not involved in vision and found no significant differences for RH7 rates between diel-niches (Fig. 2B).

Lastly, we ran the PAML analysis using a species tree instead of gene trees, since Blue and UV opsin recovery was sparse (∼50% of the taxa) and gene trees may be biased. The species tree models showed highly significant differences between diel-niches for UV, Blue and LW opsins (Fig. 2C). We did not include RH7 in this analysis in order to compare across similar species trees, which was precluded by the poor overlap of species between RH7 and visual opsins.

We tested if particular sites were under positive selection in diurnal species using branch and branch site-models. There was no significant signature of positive selection across the opsin when comparing with the null model for branch models, however this is expected for highly conserved proteins like opsins. The branch-site models, however, identified amino acid sites that were under positive selection/relaxed purifying selection in diurnal species in UV/RH4, Blue/RH5 and LW/RH6 opsins (Table S4).

Xu et al. (2013) reported elevated dN/dS (ω) rates in butterflies (diurnal) compared to moths (nocturnal)^27^, with LW, Blue and UV opsins, showing a decreasing magnitude in differences. In contrast, we find UV opsins had the highest and most consistent ω rate differences. Feuda et al. (2016) used two independent diurnal transitions, with a total of 10 Lepidoptera taxa^28^ and have results similar to ours (Fig. 2B). We found that UV/RH4 and Blue/RH5 and LW/RH6 genes in nocturnal Lepidoptera underwent strong purifying selection and more diversifying selection in diurnal species. RH7 had almost similar levels of selection in both nocturnal species and diurnal species. Only UV opsin showed consistent differences through the range of analysis parameters, therefore, we used it for 3D protein modelling.

### Mapping amino acid sites under selection to predicted protein structure

The site-selection models failed to return any significant sites under selection, but when we took diel-niche into consideration and used branch-site models, they returned several sites under positive selection (Table S4). We mapped the significant sites recovered from the branch site model onto the predicted protein structure, obtained using transmembrane helix prediction. We found that each class of opsin mapped to a unique pair of adjacent transmembrane helices, likely responsible for tuning rhodopsin (Fig. S4). We found that the for Blue and LW opsins, sites under selection, predicted by the models, were unaffected by the choice of gene vs. species trees. The choice of tree, however, affected the results of the UV opsin selection models. Since species tree models predicted more sites in the helical regions–which can affect and tune the opsin–we infer that the species tree models are more accurate. Additionally, UV 59 may be a convergent site of selection, as it has previously been identified in diel-transitions of fireflies^24^.

### Expansion of color vision through sequence tuning

Since the UV opsin recovered all 7 transmembrane domains, similar to the x-ray crystal structure of squid rhodopsin, we modelled it using an online protein modelling tool, Swiss-model^42^, and identified putative retinal binding sites, i.e. amino acids less than a minimum distance from the retinal molecule. We found that majority of the amino acid sites under positive selection are close to amino acids of the retinal binding region (Fig. 3). Feuda et al. (2016) also map the positively selected amino acids to opsins^28^, but they recover a large fraction of sites in terminal regions, not in the helices. This could be because their alignment method removes gap-filled regions, reducing the accuracy of structural prediction. We refrained from removing gap filled regions from the alignment, instead, we performed end trimming and limited the analysis to species that resulted in a gapless alignment. Our analyses show that these sites likely alter the opsin spectral tuning in diurnal species by changing the chemical environment surrounding retinal. Now that genetic manipulation in Lepidoptera is feasible using CRISPR-Cas9, one can determine the relative impact of these site-specific mutations on Lepidoptera opsin sensitivity and overall behavior.

**Figure 3:**
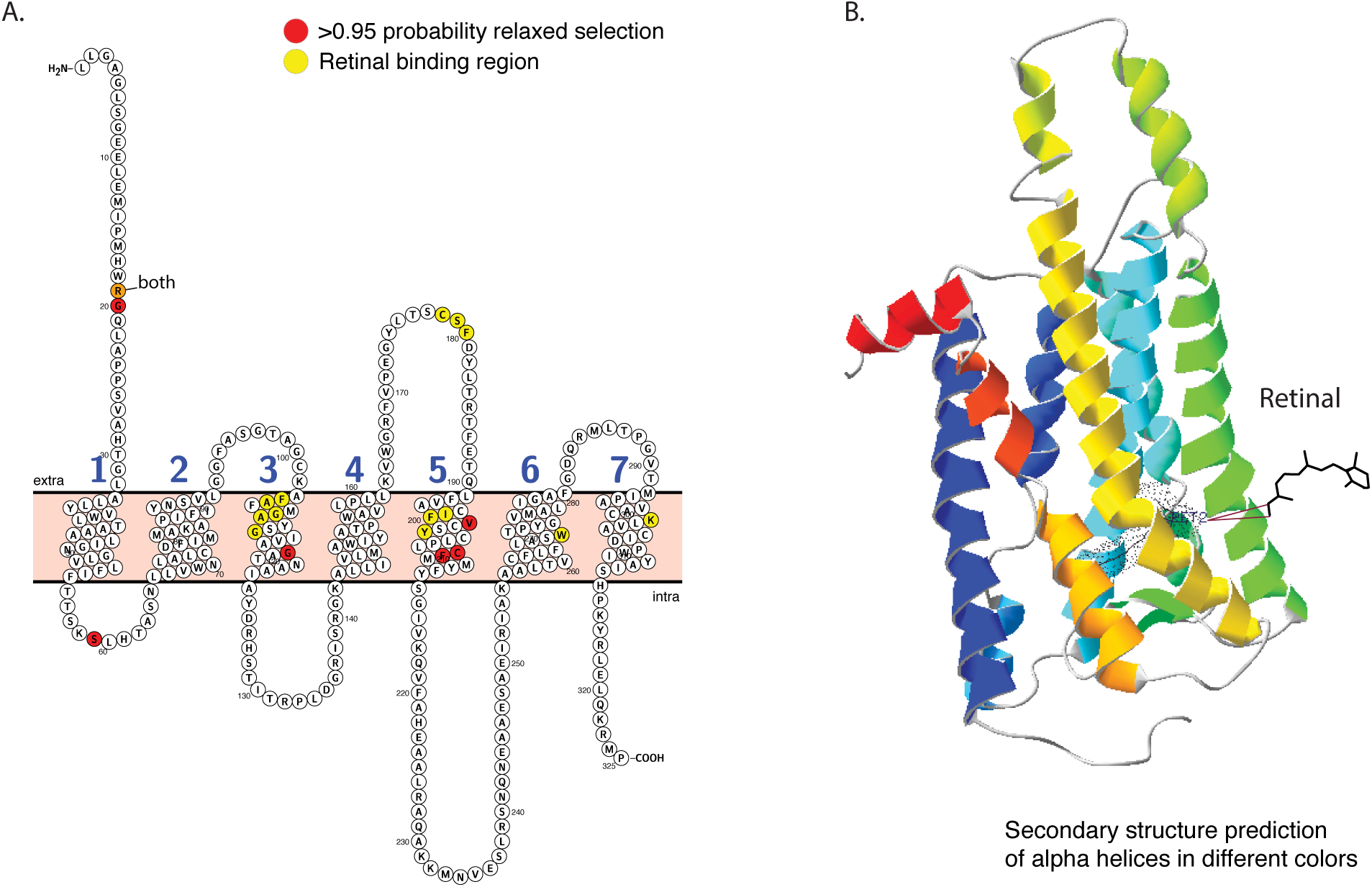
Opsin structural prediction with retinal binding sites and sites under diversifying selection. Note the overlap between a single site identified by the two models, other sites are in close proximity to each other, signifying that sites associated with rapid evolution in high light environments could be involved in opsin tuning by changing the chemical environment around retinal. A. UV-opsin PHOBIUS transmembrane prediction. Opsins sites under relaxed purifying selection and amino acids around (<=4 Å) from the retinal binding region have been marked. B. UV-opsin model made using squid rhodopsin as the template. Retinal is marked and secondary structure prediction of alpha helices are shown.

## Discussion

What does opsin expansion or contraction mean for the visual system of an organism? One consequence of opsin expansion is the potential for improved color discrimination. Color vision usually requires at least two opsins with a partial overlap in spectral sensitivity and even small changes to the amino acid sequence can modify the tune the opsin, shifting the percieved color space of an organism. Alternatively, more dramatic shifts in detectable color space can occur through loss or gain of opsins. For example many butterflies can see in the red (620-780 nm) through Long Wavelength (LW) opsin duplication and divergence^26^. RH1–6 are implicated in color vision, but non-visual RH7 is also phylogeneticly scattered across insects, and associated with circadian rhythm maintenance.

Examining opsin diversity across multiple lineages that have switched diel-niche can help determine how light environment affects opsin evolution. Dragonflies, mosquitoes and butterflies show multiple opsin duplications with as many as 15 color opsins^26,28,38,43^, while beetles and scorpionflies show losses identifying them as potential monochromats^44,45^. These studies have not examined potential links between diel activity and opsin diversity, or have failed to find consistent trends. (reviewed in^28^). Analyses are confounded by uncertainty in diel-niche assignment and a lack of multiple independent diel-switches. Systematic error — including shallow sequencing and poorly resolved species trees — could also obscure any trends ^46,47^.

The greatest limitation of large-scale gene mining approaches is the reduced power to detect absences. Low coverage transcriptomes from older studies, mixed tissue sources, as well as varied assembly and sequencing methods, all increase heterogeneity in opsin recovery. Ideally, we would sequence only eye tissue with muscle tissue from the body as a control, but we frequently lack compete information about which tissues generated the transcriptomes. Further, we do not measure expression levels, which could identify non-functional opsins^48^. However, as we verified our pipeline with known data and used multiple annotation methods and ancestral state reconstruction to identify false positives, we believe we have obtained the best estimate of opsin diversity possible from this dataset.

More than 80% of the opsin duplications we discovered across 10 independent lineages are in diurnal species, which additionally have more variation in opsin tuning sites than nocturnal species. Differences in selection rate between the diel niches are visible across all three visual opsins, but most robust and consistent in ultraviolet-sensitive opsins. Structural modelling shows that in diurnal species, amino acids opsin under positive selection overlap with the retinal binding region. Each of these observations supports our prediction that transitions from dim to bright-light environments drive opsin diversification in Lepidoptera. Only an adaptive selective force would likely show consistent patterns across multiple independent duplication events. The ancestors of both diurnal and nocturnal lepidopterans were likely crepuscular with limited trichromatic vision, so what is likely to have driven this selection?^29^

Moving to diurnal behavior reduces light constraints but increases the risk of predators, which may in turn drive selection of either chemical defenses or faster escape flights. Chemical defenses are often precursors to bright warning coloration ^49^ which can then become targets for sexual selection, which then requires accurate color perception. Aposematic *Heliconius* butterflies, for example, use wing-coloration as mate choice signals ^25,34,50^. Lepidopterans are fast in the air, with speeds up to 3.5 m s^-1^ in free flying *Danaus sp.* ^51,52^, and 5.3 m s^-1^ in *Manduca sexta* ^53^. Aposematism and fast flight appear to be alternative strategies, since most aposematic butterflies are slower than their counterparts.^52^ Butterflies and diurnal hawkmoths are both predominantly nectar feeders and the energetic requirements for fast flight may select for efficient nectar foraging; for which an expanded color space would be beneficial. In sum, the transition to a diurnal light environment may set into motion a diverse trajectory of evolutionary scenarios, each of which may increase opsin diversity.

For species that switched to a more nocturnal lifestyle, light intensity was a limiting factor, and their eyes underwent transitions to become superposition eyes; transitions that happened multiple times across Lepidoptera ^36,54^. The nature of visual pigments makes capturing color information harder as light intensity drops; stacking more rhodopsin to make more sensitive receptors comes at the cost of getting a more broadband signal and losing color discrimination ^55^. Superposition eyes effectively act as a large lens in order to increase the light available at each photoreceptor. The maintenance of trichromacy in nocturnal Lepidoptera is therefore puzzling: either color vision has some critical function in nocturnal Lepidoptera, or opsins are maintained for functions independent of color vision.

As a critical function, color can serve as a short-range cue useful for mating and foraging, such as tiger moths that can distinguish conspecifics using color markings that are unrecognizable to birds ^56^, or the strong innate attraction of flower foraging Lepidoptera to blue ^57,58^. Moths may overcome dim light constraints by pooling from different opsins. The most detailed study of opsin distribution across a moth eye shows that there are very different dorsal-ventral patterns in *Manduca* compared to butterflies^59^, suggestung that moths may have partitioned color and spatial vision in different regions.

Alternatively, selection could act on opsins independently, regardless of their utility for color vision. UV and LW opsins models consistently showed differences across models, but Blue opsin rates had a smaller differences in all but one model. LW opsin recovery was almost complete, but UV and Blue opsin recovery was patchy. If this trend represents actual losses, it supports the idea that each opsin is maintained independently and LW might be more critical. UV light sensitivity is prevalent in nocturnal animals, even dichromats such as rodents, owls and deep-sea fishes ^20,60,61^. UV contrast is a foraging cure for moths, as nectar guides in many-night blooming flowers ^62^, and an attractant ^63–65^. Because short wavelengths increase around twilight^66^, UV light is a possible signal for pupil responses, (anecdotally more prevalent across nocturnal moths than butterflies), which could explain the purifying selection in nocturnal species. LW sensitivity is useful for oviposition behavior in butterflies^67^ and their high recovery and strong purifying selection in nocturnal species could be a signature of their role in oviposition. Butterflies display strong host plant specificity compared to moths,^68–70^ and butterfly LW may have diversified for finer oviposition site discrimination. Irrespective of their role in color vision, LW and UV opsins may diversify in diurnal species but be held under purifying selection in nocturnal species due to integral functions.

In summary, patterns of opsin diversity and evolution across Lepidptera and insects show that differences in diel-niche correlate with opsin duplication and rates of evolution. Future studies on fine-scale expression patterns in closely related diel-transitions can examine opsin expression levels and immunohistochemistry to determine if the duplications are functional, and in-vitro expression systems and CRISPR could confirm that the amino acids identified tune color vision. A more thorough sampling across insect transcriptomes might identify deeper evolutionary patterns. And examining opsin distribution patterns across lepidopterans could help to understand if sensitivity and color vision trade-off in different eye regions. This study provides a library of opsin sequences, useful for these sorts of studies, as well as opsin diversity patterns across different species, which can help inform which species to probe.

## Methods

### Transcriptome annotations and database mining

#### Insect opsin gene annotation

We annotated 30 assembled insect transcriptomes from InsectBase ^30^ and Ensembl Metazoa ^71^, limiting our analyses to head or whole-body tissue to enhance recovery of visual genes (Table S3). Transcriptomes were annotated using Phylogenetically Informed Annotation (PIA) ^32^. PIA is a bioinformatic pipeline that queries transcriptomes using pre-existing reference gene-sets and places them on a supplied amino acid gene tree. It identifies reading frame and creates a comprehensive gene tree. A modified version of this pipeline was used allow for faster analysis on a high-performance cluster (https://github.com/xibalbanus/PIA2, Supplementary Information).

A set of well characterized metazoan visual rhodopsin genes were used as the reference gene set for PIA, and parameters were set to ensure high fidelity of hits while allowing for partial length matches (minimum amino acid sequence length −30, gene search type -single, gene set -r_opsin, e-value threshold -e^-19, maximum of blast hits retained −100)

We further obtained protein sequences from published genome and opsin annotations of 20 additional taxa (Table S2). These sequences were included in the opsin annotation tree (Fig. 1), but were not run through the PIA pipeline– PIA can only use nucleotide transcript data. An amino acid opsin gene tree was built to confirm the annotations (Fig. S2).

#### Lepidoptera opsin gene annotation

We annotated 162 Lepidoptera transcriptomes obtained from previously published studies (Table S1). See Kawahara et al. (2019) Supplementary Information for details on transcriptomes.^33^ PIA was used with the same parameters as described in the previous section for insects.

Further manual curation was done using the reconstructed opsin tree (Fig. S1). Putative duplications were aligned to ensure they were not fragments of the same gene. We added opsin sequences from 14 Lepidoptera species obtained from blast searches of Lepbase, a repository of lepidoptera genomes and transcriptomes,(Challis et al. 2016) and Ensembl Metazoa. We used *Manduca sexta* RH4, RH5, RH6 and, RH7 opsin sequences as queries for the BLAST search (-evalue 1.0e-10 -num_alignments 250).

#### Opsin gene tree reconstruction

The Lepidoptera and insect cDNA sequences were collected and annotated using PIA as putative short wavelength (SW/UV/RH4), medium wavelength (MW/Blue/RH5) and long wavelength (LW/RH6), RH7 like (Table S1). However, as a confirmation, we also constructed gene trees for all the sequences. These trees also allowed us to identify unknown opsins, measure divergence of duplications and, catch incorrect annotations.

#### Nucleotide gene tree

MAFFT (v7.294b)^73^ was used to align the sequences (-auto was used and MAFFT detected the sequences as nucleotide), and IQ-TREE (multi-core v1.6.12)^74^ to build a ML tree (iqtree-s alignment_name.fasta -st DNA -bb 10000 -nt AUTO -alrt 1000). The tree was color-coded based on opsin clades, and taxon names were color-coded based on the PIA annotation, to find discrepancies between the two methods. The tree was midpoint rooted using Archaeopteryx (v0.9917 beta)(https://github.com/cmzmasek/archaeopteryx-js). Using melanopsins as an outgroup did not affect the opsin annotations, so we chose to exclude them from the gene trees (Fig. S1A–D, Supplementary Information).

#### Amino acid gene tree

MAFFT (-auto was used and mafft detected the sequences as amino acid) and IQ-TREE (iqtree -s alignment_name.fasta -st AA -bb 10000 -nt AUTO -alrt 1000) were used to build an amino acid gene tree. We used it to confirm opsin protein annotations taken from annotated genomes or other studies (Feuda et al. 2016).

### Diel niche assignment

We assigned diel-niche to compare opsin evolution, but only did so for species for which we recovered at least one opsin, limiting further analysis to these taxa. Species were classified as “diurnal”, “nocturnal”, “crepuscular” (species active at dawn or dusk) or “both” (species with some activity during the day and night) (Table S1, S2). The diel-niche was assigned using published literature, natural history databases and in consultation with experts for more obscure species (Table S1, S2).

Diel-niche is often assigned based on whether an insect is attracted to light, and can be a reliable indicator of species that are strictly nocturnal and diurnal^29^. But it often fails for crepuscular species, and species that fly during the night and the day (“both”). For example, even though *Manduca sexta* is often considered crepuscular, one study has found it is almost entirely nocturnal when compared to *Hyles lineata*, which is active both during the night and day ^75^.

We used different approaches for diel-niche assignment depending on analyses. For easy tree visualization, we used only three diel states, grouping “crepuscular” and “both” into a single category (Fig. 1). For statistical analysis, however, we tried all possible grouping combinations (Table 1). Because some diel-niche assignments are uncertain, we note this ambiguity in the dataset (Table S1). For selection analysis, models only allow two groups, and we therefore categorized species into strictly “nocturnal” and “diurnal” by assigning “crepuscular” and “both” to the “diurnal” category.

### Species-tree reconstruction

We required a well-resolved species tree for Lepidoptera and insects to perform ancestral state reconstruction, opsin rate analysis, and visualization of opsin diversity. The insect species tree (Fig. 1) was built by grafting an NCBI common taxonomy onto the Misof et al. (2014) tree, pruned using Archaeopteryx. Because only a few taxa, spread across different families, were used in this tree, relationships at the family level and lower are only representative—they do not reflect current taxonomy. For the Lepidoptera species tree we pruned a well-resolved species tree from Kawahara et al. (2019) to include only species that had at least one identified opsin. We utilized the python package ETE v3 ^77^ to prune and annotate the trees. The pruned tree was modified using python scripts (Supplementary Information) to show the number of opsins in various Lepidoptera taxa (Fig. 1).

### Statistical analysis

The python scipy stats package ^78^ was used to analyze the opsin duplications and their association with diel-niche across Lepidoptera. Custom scripts (Supplementary Information) were used to filter data—including only species for which we recovered at least one opsin—and redo the tests after excluding taxa with uncertainty or varying placement of “crepuscular” and “both” diel-niches as either “nocturnal” or “diurnal” (Table 1).

### Selection Analyses

The annotated opsin dataset from Lepidoptera was used for analyzing patterns of selection acting on the opsins and comparing them across diel-niche. Each opsin family was analyzed separately, but we limited the analysis to taxa which had recovered at least one copy of all three visual opsin sequences (UV/RH4, Blue/RH5 and LW/RH6) to make the analyses comparable between genes and datasets. Because RH7 was only used as a control, we used all species from which we recovered RH7.

We used Geneious v. 10.0.9 (https://www.geneious.com) to sort and export the sequences. The sequences were manually cleaned by trimming longer sequences and removing sequences that were too short. AliView v. 1.18.1 ^79^ with Muscle v. 3.8.31 ^80^ was used for trimming and aligning. We did not remove gap-filled regions since this can bias structural prediction. Instead, we dropped sequences or trimmed edges to reduce gaps and stop codons while optimizing the length of alignment.

For RH7, the manual method resulted in a very short alignment less than half the length of the other opsins, so Prank with TranslatorX ^81^ was used for RH7 even though it had more gaps. IQ-TREE (v. 1.6.12)^74^ was used for building the gene trees from the alignments. PAML 4.9a was used to generate various models of codon evolution and estimate site-wise synonymous (α) and non-synonymous (β) rates. The likelihood ratio test was used to determine if a site has a significantly deviant β/α (w) from the neutral/null model. We used branch, site, and branch-site models to test for positive selection and relaxation in purifying selection.

Custom python scripts were used to filtered the data by number of opsins annotated, compile sequences for the filtered species, and prune the Kawahara et al. (2019) tree to create the species-tree used in PAML (Supplementary Information). We then tested whether diel-niche (“nocturnal” or “diurnal”) has influenced opsin rate evolution for the various selection models. For selection analyses, we categorized the background branches as strictly “nocturnal” and everything else as “diurnal” and as a foreground branch. We ran these analyses under different conditions, varying the number of taxa, choice of gene tree versus species trees and with different alignment methods to test for sensitivity to these parameters (Table S4; Supplementary Information).

### Ancestral state reconstruction

We used the geiger ^82^ package in R with the SYM model and 10,000 repetitions, then used ape to plot on the pruned tree. We generated 10,000 stochastic maps for each tree in SIMMAP ^83^, which is part of the R package phytools ^84^. The methods were adapted from Kawahara et al. (2018), which also mapped diel state, without prior information on opsin transition probability. We used the SYM model for each opsin class, since we did not want to assume anything about differences in rates for losses versus gains.

Stochastic character mapping is a Bayesian approach, supposedly better and more robust than other parsimony or likelihood methods because it allows changes along branches, not just tips, and makes use of data along the nodes to make predictions. It also permits the assessment of uncertainty in character history due to topology and branch lengths^85^. SIMMAP does not allow for missing or unknown data. Therefore, all tips were coded with a discrete, unordered character state, and the taxa with missing traits were pruned from the dataset, causing some discrepancies when comparing reconstructions of opsin number and diel state (Fig. S3A). We used UV/RH4, Blue/RH5, LW/RH6, total number of opsins, and diel-niche as the discrete characters for the ancestral state reconstruction.

### Protein modeling and site mapping

We mapped amino acid sites under relaxed purifying selection, from the branch-site models, to the protein structure using Protter (Omasits et al. 2014, http://wlab.ethz.ch/protter/). Protter is a web-server based tool useful in annotating a protein sequence. It uses structural prediction tools, such as Phobius ^87^ to predict transmembrane domains, and allows the user to mark a custom set of amino acids.

To obtain the retinal binding sites, we chose UV opsin because it recovered seven transmembrane helices in Protter, similar to other known invertebrate opsins. We modeled it using the online swiss model workspace(Waterhouse et al. 2018). We used a squid rhodopsin z-ray structure as a template (2z73.1), as it had a much higher identity-score than other rohdopsin structures. We used Swisspdb viewer ^88^ to fit the template and the modeled protein using magic fit, and then selected sites within 4 Å of the retinal molecule in the modeled UV-opsin.

## Acknowledgments

We thank Jessica Liberles and Joseph Ahrens for guidance with bioinformatic analyses. We thank Heather Bracken-Grissom and Jorge Perez-Moreno for discussions, advice and assistance with bioinformatics pipelines, Ravindran Palavalli-Nettimi, Jesse W. Breinholt for feedback on manuscript drafts, Scott Cinel, Harlan Gough, David Plotkin, Ryan St. Laurent, Caroline Storer, and other members of the Kawahara lab for their assistance throughout the project. We thank John P. Currea, Nick Palmero, and Carlos Ruiz, along with other members of the Theobald lab for comments and feedback on the manuscript. The authors acknowledge University of Florida Research Computing for providing computational resources and support that have contributed to the research results reported in this publication (http://researchcomputing.ufl.edu)

Funding support for the project was through NSF DEB-1557007 and NSF IOS-1750833. The Florida International University Presidential fellowship supported YS.

## References

1. Nilsson, D. E. Eye evolution and its functional basis. Vis. Neurosci. 30, 5–20 (2013).

2. Nilsson, D. E. & Arendt, D. Eye Evolution: The Blurry Beginning. Curr. Biol. 18, R1096–R1098 (2008).

3. Land, M. F. & Nilsson, D.-E. Animal Eyes. (Oxford University Press, 2012).

4. Colley, N. J. & Nilsson, D. E. Photoreception in Phytoplankton. Integr. Comp. Biol. 56, 764–775 (2016).

5. Jékely, G. Evolution of phototaxis. Philos. Trans. R. Soc. B Biol. Sci. 364, 2795–2808 (2009).

6. Porter, M. L. et al. Shedding new light on opsin evolution. Proc. R. Soc. B Biol. Sci. 279, 3–14 (2012).

7. Henze, M. J. & Oakley, T. H. The dynamic evolutionary history of pancrustacean eyes and opsins. Integr. Comp. Biol. 55, 830–842 (2015).

8. Colbourne, J. K. et al. The Ecoresponsive Genome of *Daphnia pulex*. Science 331, 555–561 (2011).

9. Futahashi, R. et al. Extraordinary diversity of visual opsin genes in dragonflies. Proc. Natl. Acad. Sci. 112, E1247–E1256 (2015).

10. Porter, M. L. et al. The evolution of complexity in the visual systems of stomatopods: Insights from transcriptomics. Integr. Comp. Biol. 53, 39–49 (2013).

11. Marshall, J. & Arikawa, K. Unconventional colour vision. Curr. Biol. 24, R1150–R1154 (2014).

12. Barlow, H. B. What causes trichromacy? A theoretical analysis using comb-filtered spectra. Vision Res. 22, 635–643 (1982).

13. Hoffmann, M. et al. Opsin gene duplication and diversification in the guppy, a model for sexual selection. Proc. R. Soc. B Biol. Sci. (2007) doi: 10.1098/rspb.2006.3707.

14. Everett, A., Tong, X., Briscoe, A. D. & Monteiro, A. Phenotypic plasticity in opsin expression in a butterfly compound eye complements sex role reversal. BMC Evol. Biol. 12, 232 (2012).

15. Dyer, A. G. Discrimination of flower colours in natural settings by the bumblebee species *Bombus terrestris* (Hymenoptera: Apidae). Entomol. Gen. (2005) doi: 10.1127/entom.gen/28/2006/257.

16. Tierney, S. M. et al. Consequences of evolutionary transitions in changing photic environments. Austral Entomol. 56, 23–46 (2017).

17. Warrant, E. J. & Johnsen, S. Vision and the light environment. Curr. Biol. 23, R990–R994 (2013).

18. Zhao, H. et al. The evolution of color vision in nocturnal mammals. Proc. Natl. Acad. Sci. 106, 8980–8985 (2009).

19. Stern, D. B. & Crandall, K. A. Phototransduction Gene Expression and Evolution in Cave and Surface Crayfishes. Integr. Comp. Biol. 58, 398–410 (2018).

20. Musilova, Z. et al. Vision using multiple distinct rod opsins in deep-sea fishes. Science 364, 588–592 (2019).

21. Tobler, M., Coleman, S. W., Perkins, B. D. & Rosenthal, G. G. Reduced opsin gene expression in a cave-dwelling fish. Biol. Lett. 6, 98–101 (2010).

22. Schweikert, L. E., Fitak, R. R., Caves, E. M., Sutton, T. T. & Johnsen, S. Spectral sensitivity in ray-finned fishes: Diversity, ecology and shared descent. J. Exp. Biol. 221, (2018).

23. Macias-Muñoz, A., Rangel Olguin, A. G. & Briscoe, A. D. Evolution of Phototransduction Genes in Lepidoptera. Genome Biol. Evol. 11, 2107–2124 (2019).

24. Sander, S. E. & Hall, D. W. Variation in opsin genes correlates with signalling ecology in North American fireflies. Mol. Ecol. (2015) doi: 10.1111/mec.13346.

25. Briscoe, A. D. Reconstructing the ancestral butterfly eye: focus on the opsins. J. Exp. Biol. 211, 1805–1813 (2008).

26. Arikawa, K. The eyes and vision of butterflies. J. Physiol. 595, 5457–5464 (2017).

27. Xu, P. et al. The Evolution and Expression of the Moth Visual Opsin Family. PLoS One 8, e78140 (2013).

28. Feuda, R., Marlétaz, F., Bentley, M. A. & Holland, P. W. H. Conservation, duplication, and divergence of five opsin genes in insect evolution. Genome Biol. Evol. 8, 579–587 (2016).

29. Kawahara, A. Y. et al. Diel behavior in moths and butterflies: a synthesis of data illuminates the evolution of temporal activity. Organisms Diversity and Evolution vol. 18 13–27 (2018).

30. Yin, C. et al. InsectBase: a resource for insect genomes and transcriptomes. Nucleic Acids Res. 44, D801–D807 (2016).

31. Kersey, P. J. et al. Ensembl Genomes 2018: an integrated omics infrastructure for non-vertebrate species. Nucleic Acids Res. 46, D802–D808 (2018).

32. Speiser, D. I. et al. Using phylogenetically-informed annotation (PIA) to search for light-interacting genes in transcriptomes from non-model organisms. BMC Bioinformatics (2014) doi: 10.1186/s12859-014-0350-x.

33. Kawahara, A. Y. et al. Phylogenomics reveals the evolutionary timing and pattern of butterflies and moths. Proc. Natl. Acad. Sci. 116, 22657–22663 (2019).

34. Briscoe, A. D. et al. Positive selection of a duplicated UV-sensitive visual pigment coincides with wing pigment evolution in *Heliconius* butterflies. Proc. Natl. Acad. Sci. U. S. A. 107, 3628–33 (2010).

35. Andersson, S., Rydell, J. & Svensson, M. G. E. Light, predation and the lekking behaviour of the ghost swift *Hepialus humuli* (L.) (Lepidoptera, Hepialidae). Proc. R. Soc. London. Ser. B Biol. Sci. 265, 1345–1351 (1998).

36. Pirih, P. et al. The giant butterfly-moth *Paysandisia archon* has spectrally rich apposition eyes with unique light-dependent photoreceptor dynamics. J. Comp. Physiol. A 204, 639–651 (2018).

37. Kinoshita, M. & Arikawa, K. Color and polarization vision in foraging *Papilio*. J. Comp. Physiol. A Neuroethol. Sensory, Neural, Behav. Physiol. 200, 513–526 (2014).

38. Chen, P. J., Awata, H., Matsushita, A., Yang, E.-C. & Arikawa, K. Extreme spectral richness in the eye of the common bluebottle butterfly, Graphium sarpedon. Front. Ecol. Evol. 4, (2016).

39. Langer, H., Hamann, B. & Meinecke, C. Tetrachromatic Visual System in the Moth Spodoptera exempta. J. Comp. Physiol. A 129, 235–239 (1979).

40. Fu, X., Feng, H., Liu, Z. & Wu, K. Trans-regional migration of the beet armyworm, *Spodoptera exigua* (Lepidoptera: Noctuidae), in North-East Asia. PLoS One 12, e0183582 (2017).

41. Telles, F. J. et al. Out of the blue: The spectral sensitivity of hummingbird hawkmoths. J. Comp. Physiol. A Neuroethol. Sensory, Neural, Behav. Physiol. 200, 537–546 (2014).

42. Waterhouse, A. et al. SWISS-MODEL: homology modelling of protein structures and complexes. Nucleic Acids Res. 46, W296–W303 (2018).

43. Bybee, S. M., Johnson, K. K., Gering, E. J., Whiting, M. F. & Crandall, K. A. All the better to see you with: A review of odonate color vision with transcriptomic insight into the odonate eye. Org. Divers. Evol. 12, 241–250 (2012).

44. Böhm, A., Meusemann, K., Misof, B. & Pass, G. Hypothesis on monochromatic vision in scorpionflies questioned by new transcriptomic data. Sci. Rep. 8, 1–8 (2018).

45. Lord, N. P. et al. A cure for the blues: Opsin duplication and subfunctionalization for short-wavelength sensitivity in jewel beetles (Coleoptera: Buprestidae). BMC Evol. Biol. 16, (2016).

46. Yang, Z. PAML 4: Phylogenetic Analysis by Maximum Likelihood. Mol. Biol. Evol. 24, 1586–1591 (2007).

47. Diekmann, Y. & Pereira-Leal, J. B. Gene Tree Affects Inference of Sites under Selection by the Branch-Site Test of Positive Selection. Evol. Bioinforma. 11s2, EBO.S30902 (2015).

48. Wagner, G. P., Kin, K. & Lynch, V. J. A model based criterion for gene expression calls using RNA-seq data. Theory Biosci. 132, 159–164 (2013).

49. Sherratt, T. N. The coevolution of warning signals. Proc. R. Soc. London. Ser. B Biol. Sci. 269, 741–746 (2002).

50. Morehouse, N. I. & Rutowski, R. L. In the Eyes of the Beholders: Female Choice and Avian Predation Risk Associated with an Exaggerated Male Butterfly Color. Am. Nat. 176, 768–784 (2010).

51. Dudley & Srygley. Flight physiology of Neotropical butterflies: allometry of airspeeds during natural free flight. J. Exp. Biol. 191, 125–39 (1994).

52. Dudley, R. Biomechanics of flight in neotropical butterflies: Morphometrics and kinematics. J. Exp. Biol. 150, 37–53 (1990).

53. Stevenson, R., Corbo, K., Baca, L. & Le, Q. Cage size and flight speed of the tobacco hawkmoth *Manduca sexta*. J. Exp. Biol. 198, 1665–72 (1995).

54. Nilsson, D. E., Land, M. F. & Howard, J. Optics of the butterfly eye. J. Comp. Physiol. A 162, 341–366 (1988).

55. Cronin, T. W., Johnsen, S., Marshall, N. J. & Warrant, E. J. Visual Ecology. Visual Ecology (Princeton University Press, 2014). doi: 10.23943/princeton/9780691151847.001.0001.

56. Henze, M. J., Lind, O., Mappes, J., Rojas, B. & Kelber, A. An aposematic colour-polymorphic moth seen through the eyes of conspecifics and predators – Sensitivity and colour discrimination in a tiger moth. Funct. Ecol. 32, 1797–1809 (2018).

57. Kelber, A. Innate preferences for flower features in the hawkmoth *Macroglossum stellatarum*. J. Exp. Biol. 200, 827–36 (1997).

58. Yurtsever, S., Okyar, Z. & Guler, N. What colour of flowers do Lepidoptera prefer for foraging? Biologia (Bratisl). 65, 1049–1056 (2010).

59. White, R. H. The retina of Manduca sexta: rhodopsin expression, the mosaic of green-, blue- and UV-sensitive photoreceptors, and regional specialization. J. Exp. Biol. 206, 3337–3348 (2003).

60. Jacobs, G. H., Fenwick, J. A. & Williams, G. A. Cone-based vision of rats for ultraviolet and visible lights. J. Exp. Biol. 204, 2439–46 (2001).

61. Höglund, J. et al. Owls lack UV-sensitive cone opsin and red oil droplets, but see UV light at night: Retinal transcriptomes and ocular media transmittance. Vision Res. 158, 109–119 (2019).

62. Hirota, S. K., Miki, N., Yasumoto, A. A. & Yahara, T. UV bullseye contrast of *Hemerocallis* flowers attracts hawkmoths but not swallowtail butterflies. Ecol. Evol. 9, 52–64 (2019).

63. Robinson, H. S. & Robinson, P. J. Some notes on the observed behaviour of Lepidoptera ln flight in the vicinity of light sources. Entomol. Gaz. 1, 3–15 (1950).

64. Lamarre, G. P. A. et al. Stay Out (Almost) All Night: Contrasting Responses in Flight Activity Among Tropical Moth Assemblages. Neotrop. Entomol. 44, 109–115 (2015).

65. van Langevelde, F., Ettema, J. A., Donners, M., WallisDeVries, M. F. & Groenendijk, D. Effect of spectral composition of artificial light on the attraction of moths. Biol. Conserv. 144, 2274–2281 (2011).

66. Johnsen, S. et al. Crepuscular and nocturnal illumination and its effects on color perception by the nocturnal hawkmoth Deilephila elpenor. J. Exp. Biol. 209, 789–800 (2006).

67. Kelber, A. Ovipositing butterflies use a red receptor to see green. J. Exp. Biol. 202, 2619–2630 (1999).

68. Ehrlich, P. R. & Raven, P. H. Butterflies and Plants: A Study in Coevolution. Evolution (N. Y). 18, 586–608 (1964).

69. Holm, S. et al. No Indication of High Host-Plant Specificity in Afrotropical Geometrid Moths. J. Insect Sci. 19, (2019).

70. Scoble, M. The Lepidoptera. Form, function and diversity. (Oxford University Press, 1992).

71. Kersey, P. J. et al. Ensembl Genomes 2018: an integrated omics infrastructure for non-vertebrate species. Nucleic Acids Res. (2017) doi: 10.1093/nar/gkx1011.

72. Challis, R. J., Kumar, S., Dasmahapatra, K. K. K., Jiggins, C. D. & Blaxter, M. Lepbase: the Lepidopteran genome database. bioRxiv 056994 (2016) doi: 10.1101/056994.

73. Katoh, K. & Standley, D. M. MAFFT Multiple Sequence Alignment Software Version 7: Improvements in Performance and Usability. Mol. Biol. Evol. 30, 772–780 (2013).

74. Trifinopoulos, J., Nguyen, L.-T., von Haeseler, A. & Minh, B. Q. W-IQ-TREE: a fast online phylogenetic tool for maximum likelihood analysis. Nucleic Acids Res. 44, W232–W235 (2016).

75. Broadhead, G. T., Basu, T., von Arx, M. & Raguso, R. A. Diel rhythms and sex differences in the locomotor activity of hawkmoths. J. Exp. Biol. 220, 1472–1480 (2017).

76. Misof, B. et al. Phylogenomics resolves the timing and pattern of insect evolution. Science 346, 763–767 (2014).

77. Huerta-Cepas, J., Serra, F. & Bork, P. ETE 3: Reconstruction, Analysis, and Visualization of Phylogenomic Data. Mol. Biol. Evol. 33, 1635–1638 (2016).

78. Virtanen, P. et al. SciPy 1.0--Fundamental Algorithms for Scientific Computing in Python. (2019).

79. Larsson, A. AliView: a fast and lightweight alignment viewer and editor for large datasets. Bioinformatics 30, 3276–3278 (2014).

80. Edgar, R. C. MUSCLE: multiple sequence alignment with high accuracy and high throughput. Nucleic Acids Res. 32, 1792–1797 (2004).

81. Abascal, F., Zardoya, R. & Telford, M. J. TranslatorX: Multiple alignment of nucleotide sequences guided by amino acid translations. Nucleic Acids Res. 38, W7–W13 (2010).

82. Pennell, M. W. et al. Geiger v2.0: An expanded suite of methods for fitting macroevolutionary models to phylogenetic trees. Bioinformatics (2014) doi: 10.1093/bioinformatics/btu181.

83. Bollback, J. P. SIMMAP: Stochastic character mapping of discrete traits on phylogenies. BMC Bioinformatics 7, (2006).

84. Revell, L. J. phytools: an R package for phylogenetic comparative biology (and other things). Methods Ecol. Evol. 3, 217–223 (2012).

85. Revell, L. J. Two new graphical methods for mapping trait evolution on phylogenies. Methods Ecol. Evol. 4, 754–759 (2013).

86. Omasits, U., Ahrens, C. H., Müller, S. & Wollscheid, B. Protter: Interactive protein feature visualization and integration with experimental proteomic data. Bioinformatics (2014) doi: 10.1093/bioinformatics/btt607.

87. Kall, L., Krogh, A. & Sonnhammer, E. L. L. Advantages of combined transmembrane topology and signal peptide prediction--the Phobius web server. Nucleic Acids Res. 35, W429–W432 (2007).

88. Guex, N., Peitsch, M. C. & Schwede, T. Automated comparative protein structure modeling with SWISS-MODEL and Swiss-PdbViewer: A historical perspective. Electrophoresis 30, S162–S173 (2009).

